# Rethinking Niche Conservatism with Phylogenetic Location-Scale Models

**DOI:** 10.1101/2025.03.13.643081

**Authors:** Ben Halliwell

## Abstract

Phylogenetic niche conservatism (PNC) describes the tendency of evolutionary lineages to retain functional traits over time, leading to closely related species exhibiting similar phenotypes and ecological niches. Traditional approaches to studying PNC focus on phylogenetic signal in trait means, often overlooking the conservation of trait variability across lineages. This oversight limits our understanding of the ecological constraints on species adaptation. Trait variance, influenced by genetic, ecological, and environmental factors, is crucial for understanding evolutionary processes and species’ adaptive potential. Phylogenetic location-scale models offer a promising framework to incorporate both mean and variance components of trait evolution, providing a more nuanced understanding of how ecological and evolutionary processes shape trait evolution across lineages. We apply this approach to leaf economic data for the genus Eucalyptus, demonstrating that models including predictors for both mean and variance offer greater predictive performance and deeper insights into the mechanisms of niche conservatism and ecological adaptation. Our findings highlight the importance of considering trait variance in studies of PNC and its implications for assessing species’ resilience to environmental change.

## 1 Introduction

Phylogenetic niche conservatism (PNC) describes the tendency of evolutionary lineages to retain functional traits over time, leading to closely related species exhibiting similar phenotypes and ecological niches (Wiens et al., 2010). This phenomenon has profound implications for understanding ecological and evolutionary processes, including species distribution patterns, community assembly, biodiversity, ecosystem function and resilience to environmental change (Crisp and Cook, 2012; Crisp et al., 2009). Traditional approaches to studying PNC focus exclusively on phylogenetic signal in trait means, typically measured through statistics such as Blomberg’s *K* Blomberg et al. (2003) or Pagel’s *λ* (Pagel, 1999). While this approach has provided valuable insights, it overlooks a critical aspect of eco-evolutionary processes: the conservation of trait variability across lineages. This incomplete view of trait conservatism potentially limits our understanding of the ecological constraints on species adaptation.

Trait variance, or the variability in trait values within and among species, is the raw material upon which selection acts, providing insight into evolutionary processes. Ecological specialization, driven by directional and stabilizing selection, often leads to reductions in trait variance as species adapt to specific environmental conditions or niches. Under strong abiotic constraints—such as extreme temperatures or limited resource availability—species may evolve narrow physiological tolerances, resulting in reduced variability in functional traits. Biotic interactions, such as competition or predation, further constrain trait variability, as species adapt to specific ecological roles within communities.

A quantitative understanding of between-species variation in trait variability would be valuable for many applied goals, such as assessing species vulnerability to environmental change, invasive-ness, and the breeding potential of wild crop relatives. However, drivers of trait variance have largely been ignored in studies of PNC, as in biology more broadly (Cleasby and Nakagawa, 2011; Kneib et al., 2023). This oversight is particularly striking given that trait variance is often influenced by the same forces that shape mean trait values, suggesting the potential for mean-variance relationships. For instance, adaptation by directional and stabilizing selection not only maintains mean trait values within a certain range but also reduces trait variance around the mean, leading to greater phenotypic uniformity, even canalization of trait values, within different lineages. Conversely, divergent selection or ecological release can increase trait variance, allowing lineages to explore new niches or adapt to novel environments, with implications for species resilience to rapid environmental change (Crisp et al., 2009).

Phylogenetic location-scale models offer a promising framework for redefining PNC to incorporate both mean and variance components of trait evolution (Nakagawa et al., 2025). These models, which simultaneously estimate phylogenetic signal in both the location (mean) and scale (variance) of trait distributions, provide a more nuanced understanding of how ecological and evolutionary processes shape trait evolution across lineages. By explicitly modeling trait variance, location-scale models can reveal patterns of conserved variability that may be missed by traditional approaches. For example, closely related species may exhibit similar levels of trait variability due to shared evolutionary constraints or ecological pressures, even if their mean trait values diverge. Like the mean, trait variance can be influenced by genetic, ecological or environmental factors, and may reflect sampling hierarchies in the data (O’Dea et al., 2022; Westneat et al., 2015). Location-scale models explore this by allowing separate model formulas to be specified for each of these parameters. For instance, fixed and random predictors can be applied to both the mean and variance terms, allowing for the inclusion of covariates, random effects that account for grouping or phylogenetic relatedness, as well as correlations between random effects in the mean and variance terms. This dual focus on mean and variance offers deeper insights into the mechanisms of niche conservatism and ecological adaptation, and a robust framework for investigating the adaptive potential of species in changing environments. Moreover, flexibly specifying predictors for the mean and variance has the potential to enhance model predictive performance, including imputation of missing data.

We apply this approach to leaf economic data for the genus Eucalyptus to explore applications to common species functional traits. Phylogenetic location-scale models support fixed environmental predictors in both the mean and variance of LMA. We further detect phylogenetic signal in both of these parameters, and find evidence of a between-species mean-variance relationship. Location-scale models also provided greater predictive performance based leave-one-out (LOO) cross validation (CV).

## 2 Model

A general notation for linear mixed models is given by,

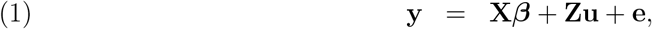

where **y** is a vector of observations. The design matrices **X** and **Z** relate fixed and random predictors to the data, the corresponding parameter vectors ***β*** and **u** contain the fixed and random effects to be estimated, and **e** is a vector of residual errors. The same notation can be adopted for a multiple response model, where 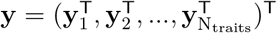 contains observations for all traits, stacked in a single column vector. To illustrate, we represent the linear predictor for the bivariate case by placing each trait in a separate row,

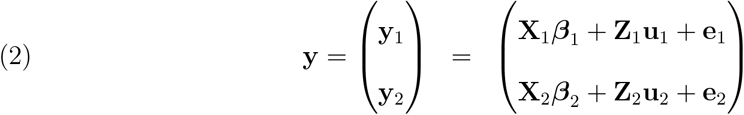

with

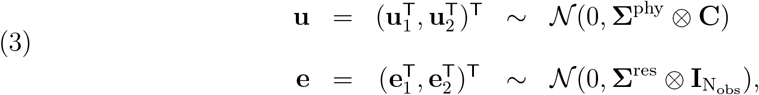

where **X**_1_***β***_1_ and **Z**_1_**u**_1_ are the linear predictors for the fixed and random effects, respectively, for response trait **y**_1_, and similarly for **y**_2_.

### 2.1 Distributional Models

Distributional models allow separate prediction for each parameter of the response distribution, e.g., the mean and variance of a Gaussian distribution. Multilevel distributional models that include random intercepts in both the mean and variance, for a common grouping variable, introduce covariance and heteroskedasticity (non-constant variance), respectively.

To illustrate, consider the MR-PMM in (2). **u** is a *location* random effect, adding a normally distributed random intercept to the model for the *mean*, which captures covariance attributable to the grouping variable. However, the model remains homoskedastic across species—all observations, irrespective of their phylogenetic effects, share a common residual variance—unless the model for the residual variance also depends on the group structure. To implement a group-structured variance model, we can specify

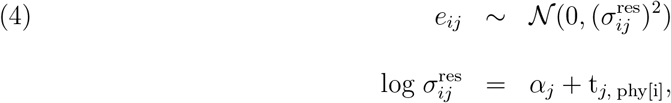

where, 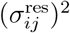 is the residual variance of the *i*^*th*^ observation for the *j*^*th*^ trait, *α*_*j*_ is the fixed intercept for the variance of the *j*^*th*^ trait, and t_*j*, phy[i]_ is a trait-specific random phylogenetic effect. To include the possibility of correlations between the random phylogenetic effects appearing in both the mean and variance, and thus share information across the distributional parameters, it is preferable to specify a joint prior distribution (Bürkner, 2019),

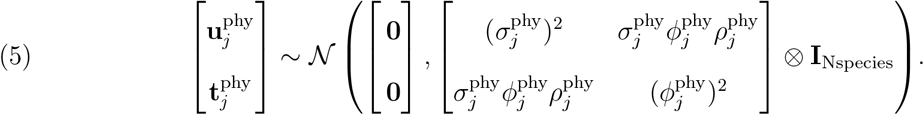

While location-scale models are increasingly used to understand variation in intra-individual behavioural variability (Mitchell et al., 2021; O’Dea et al., 2022), models such as (5) appear to be severely underutilised in phylogenetic comparative analyses (Horvath et al., 2023; Cleasby and Nakagawa, 2011). Indeed, the mean and variance of biological traits are often positively correlated (a special case of heteroskedasticity), routinely threatening the assumption of homogeneous variances in common statistical tests. In some instances, this issue is addressed by the mean-variance relationship determined by the choice of statistical distribution and link function (e.g., Poisson with log link), or else using variance-stabilising transformations (e.g., log transforming response data). However, heteroskedasticity may remain in spite of these efforts, and this ‘variance of the variance’ may in fact correspond to interesting biological effects that we would like to model explicitly. As with our model for the mean, we may include an arbitrary number of predictors for the variance, including fixed effects (e.g., methodological or environmental covariates), and random effects that contain further hierarchical structure. For example, a reasonable assumption for many phylogenetic comparative analyses is that residual variances should differ between species and that variation in residual variance should itself contain phylogenetic structure (i.e., closely related species should show similar amounts of residual variability). Furthermore, estimating correlations between phylogenetic random effects (as in 5) provides a method to test for conserved mean-variance relationships within and between traits.

For studies attempting to understand drivers of phenotypic variability, explicitly partitioning trait variance across known hierarchical structures has the potential to be extremely informative (O’Dea et al., 2022; McNeish, 2021; Westneat et al., 2015). Location-scale models also have a clear statistical motivation; heteroskedasticity violates, by definition, the assumption of homogeneous variances, leading to inefficient parameter estimation and biased errors, with potential consequences for inference and hypothesis tests.

## 3 Methods

We applied phylogenetic location-scale models to examine drivers of within- and between-species variation in a dataset of *>*6800 observations of leaf mass per area (LMA) across 685 species of Eucalyptus. The data had a complex hierarchical structure, with observations nested within species and studies. We asked whether models including fixed and random predictors for the variance provided more nuanced inferences, or better predictive performance, over models restricted to the mean. It is important to emphasize that the formula supplied for the scale term relates to the residual variance of the model, which is conditional on any fixed and random predictors included for the mean. Careful consideration must therefore be given to model structure when interpreting these effects. For an intercept only model, residuals represent deviations from the population trait mean. For a model including a fixed continuous predictor, the residuals represent deviations from the fitted regression line. For a model including a random intercept for a grouping variable (e.g., species), the residuals represent deviations from the conditional group mean, after accounting for any fixed effects.

We first fit a series of models specifying different combinations of grouping variables in both the mean and variance terms, capturing phylogenetic between-species, non-phylogenetic between-species, and between-study effects. When excluding fixed effects, these models allow us to evaluate the strength of phylogenetic signal in both the mean and variance of LMA. Including fixed effects further allows us to test the influence of covariates on both the mean and residual variance terms, and evaluate the importance of grouping variables conditional on these covariates. We compared the predictive performance of models including and excluding phylogenetic random effects in the mean and variance terms, as well as correlations between random effects in the mean and variance terms, using leave-one-out cross validation (CV). All models were fit in R version 4.3.2 using the brms package.

## 4 Results

Despite strong phylogenetic signal in the mean of LMA, we found limited evidence for phylogenetic signal in the residual variance of LMA (Figure 1A,B,C). Non-phylogenetic between-species effects revealed that species nonetheless showed consistent differences in the extent of LMA variability (Figure 1B). Furthermore, a negative correlation between non-phylogenetic between-species random effects in the mean and the variance indicated that Eucalyptus species with higher mean LMA also show lower variability in LMA (Figure 1D). The correlation between phylogenetic random effects in the mean and variance was also negative, but highly uncertain. Importantly, models that included predictors for the variance vastly outperformed models restricted to the mean according to LOO-CV (Figure 2). The preferred model (the simplest model with a loo score within 1 se error of the best performing model Yates et al. (2022)) included all group effects in the model for the mean (phylogenetic, non-phylogenetic and study), but only non-phylogenetic and study effects in the model for the variance.

**Figure 1:**
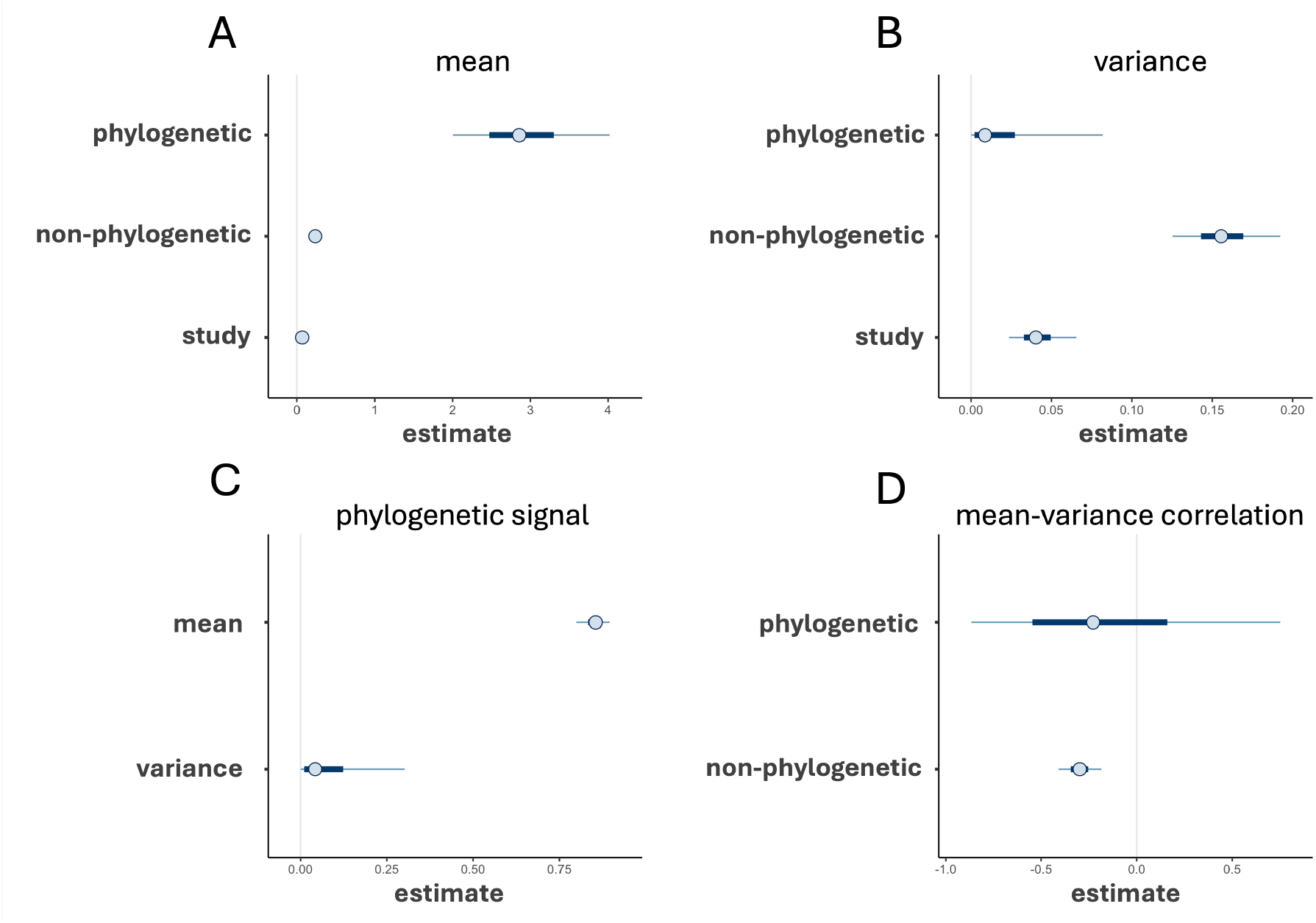
Parameter estimates from a phylogenetic location-scale model fit to LMA data from 685 Eucalyptus species. A) variance components for the modeled mean, B) variance components for the modeled variance, C) phylogenetic signal in the modeled mean and variance of LMA values, D) multilevel correlations between random effects in the mean and variance.

**Figure 2:**
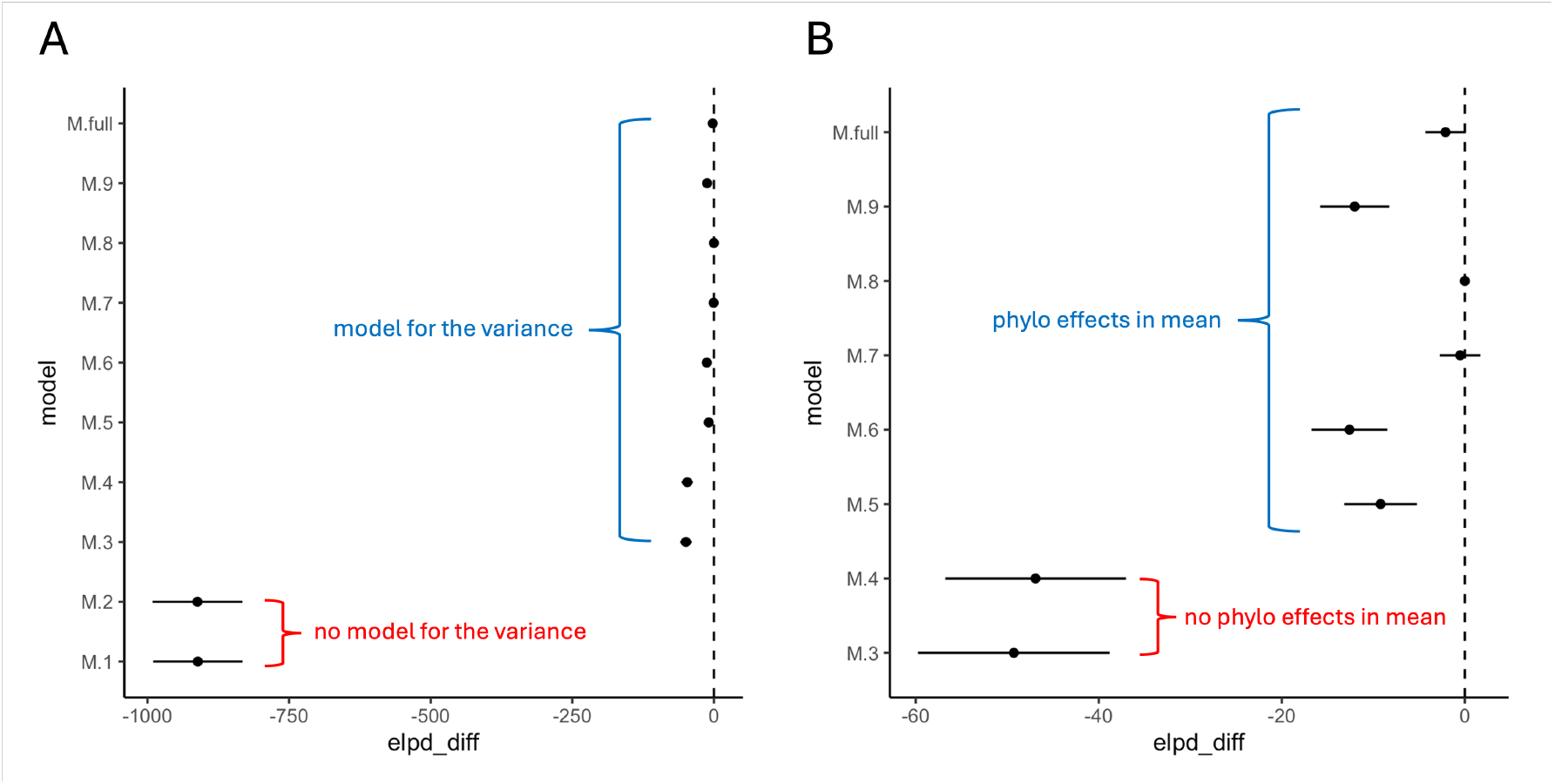
Model comparison by leave-one-out (LOO) cross validation (CV). Plots show the difference in expected log predictive density between the best performing model and other candidate models. A) Models including predictors for the variance are strongly preferred over those only including predictors for the mean, B) models including phylogenetic effects in the mean are strongly preferred over those that do not.

## 5 Discussion

PNC encompasses two distinct but interrelated concepts. First, it posits that species tend to compete effectively in environments to which they are already well suited, causing species ecology to evolve slowly, and closely related species to exhibit similar phenotypes and niches. This is evidenced by the presence of phylogenetic signal in species mean functional trait values. Second, PNC implies that ecological niches demand specific values of traits for species to persist and remain competitive. Critical traits may show limited variability under certain environmental conditions because specific values are essential for fitness. This phenomena is likely to be more pronounced within-than between-clades, where phylogenetic noise in trait-environment relationships is reduced (Anderegg, 2023). The combination of conserved trait means and strong stabilizing selection on trait values could result in conserved relationships between trait variability and environmental gradients. This dual aspect of PNC underscores the importance of considering both the evolutionary history of niche traits and the ecological constraints that shape trait variability within niches. Phylogenetic location-scale models open a a path to assess both dimensions of PNC.

From a practical perspective, phylogenetic location-scale models may offer useful inferences for forest and conservation management. Trait variance can serve as an indicator of a species’ adaptive potential or ecological flexibility, providing insights into how species may respond to environmental change or novel biotic interactions. Species with low trait variance may be more vulnerable to environmental perturbations, as they have limited capacity to adapt to new conditions. In contrast, species with high trait variance may exhibit greater resilience, adaptive potential, or invasive-ness, enabling them to persist in changing environments. We demonstrate the potential for dramatic increases in predictive performance from location-scale models, highlighting benefits for phylogenetic imputation.

## 6 Acknowledgements

We thank Luke A. Yates, Barbara R. Holland and Mark Westoby for discussions that inspired this manuscript.

